# Neural circuit basis of aversive odour processing in *Drosophila* from sensory input to descending output

**DOI:** 10.1101/394403

**Authors:** Paavo Huoviala, Michael-John Dolan, Fiona M. Love, Philip Myers, Shahar Frechter, Shigehiro Namiki, Lovisa Pettersson, Ruairí J.V. Roberts, Robert Turnbull, Zane Mitrevica, Patrick Breads, Philipp Schlegel, Alexander Shakeel Bates, Tiago Rodrigues, Yoshinori Aso, Davi Bock, Gerald M. Rubin, Marcus Stensmyr, Gwyneth Card, Marta Costa, Gregory S.X.E. Jefferis

## Abstract

Evolution has shaped nervous systems to produce stereotyped behavioural responses to ethologically relevant stimuli. For example when laying eggs, female Drosophila avoid geosmin, an odorant produced by toxic moulds. Here we identify second, third, and fourth order neurons required for this innate olfactory aversion. Connectomics data place these neurons in a complete synaptic circuit from sensory input to descending output. We find multiple levels of valence-specific convergence, including a novel form of axo-axonic input onto second order neurons conveying another danger signal, the pheromone of parasitoid wasps. However, we also observe extensive divergence: second order geosmin neurons connect with a diverse array of 80 third order cell types. We find a pattern of convergence of aversive odour channels at this level. Crossing one more synaptic layer, we identified descending neurons critical for egg-laying aversion. Our data suggest a transition from a labelled line organisation in the periphery to a highly distributed central brain representation that is then coupled to distinct descending pathways.

---

Microbes and parasites are a major driving force of natural selection in animals. As immunological defence is costly and reactive, it is better to avoid sources of infection whenever possible (1). *Drosophila* avoid laying eggs on food contaminated with harmful moulds. This avoidance is triggered by a volatile molecule, geosmin, sensed through a single olfactory receptor (Or56a) expressed in only ~ 25 sensory neurons (ORNs) per antenna (2, 3). The olfactory system, in both vertebrates and invertebrates, is a particularly shallow sensory modality where the sensory periphery is only two synapses away from higher brain areas important for organizing behaviour and forming memories (4). The clear behavioural significance of geosmin, together with Or56a ORNs being its sole dedicated sensor, make this ‘labelled line’ pathway an attractive target for studies of how sensory signals are transformed into innate, ethologically appropriate behavioural responses. Since the challenge of differentiating suitable from contaminated food substrates is ubiquitous and encompasses many issues in sensory processing, general principles are likely to emerge from these studies.

Building on the identification of Or56a sensory neurons (2), we traced the geosmin pathway deeper into the brain. Wild-type flies avoid geosmin in an egg-laying two-choice assay, and this avoidance is solely due to olfaction via the Or56a ORNs (2) (Figure S1, A and B). However, geosmin did not decrease egg-laying quantity in a no-choice situation (Figure S1C), suggesting that the phenotype arises from positional aversion. Or56a ORNs synapse onto uniglomerular DA2 projection neurons (PNs) in the antennal lobe. We identified a sparse driver line (R85E04-GAL4, Figure 1A (5)) that potentially labelled DA2 PNs. Driving the Halo-tag reporter (6) with Or56a-GAL4 and R85E04 verified that both sets of neurons target the same glomerulus (Figure S1D) while immunostaining confirmed that the PNs are cholinergic and excitatory (Figure S1F). *In vivo* electrophysiology revealed strong and highly selective responses to geosmin (Figure S1E). ‘Labelled-line’ encoding of geosmin is therefore retained at the second order PN level.

**Figure 1:**
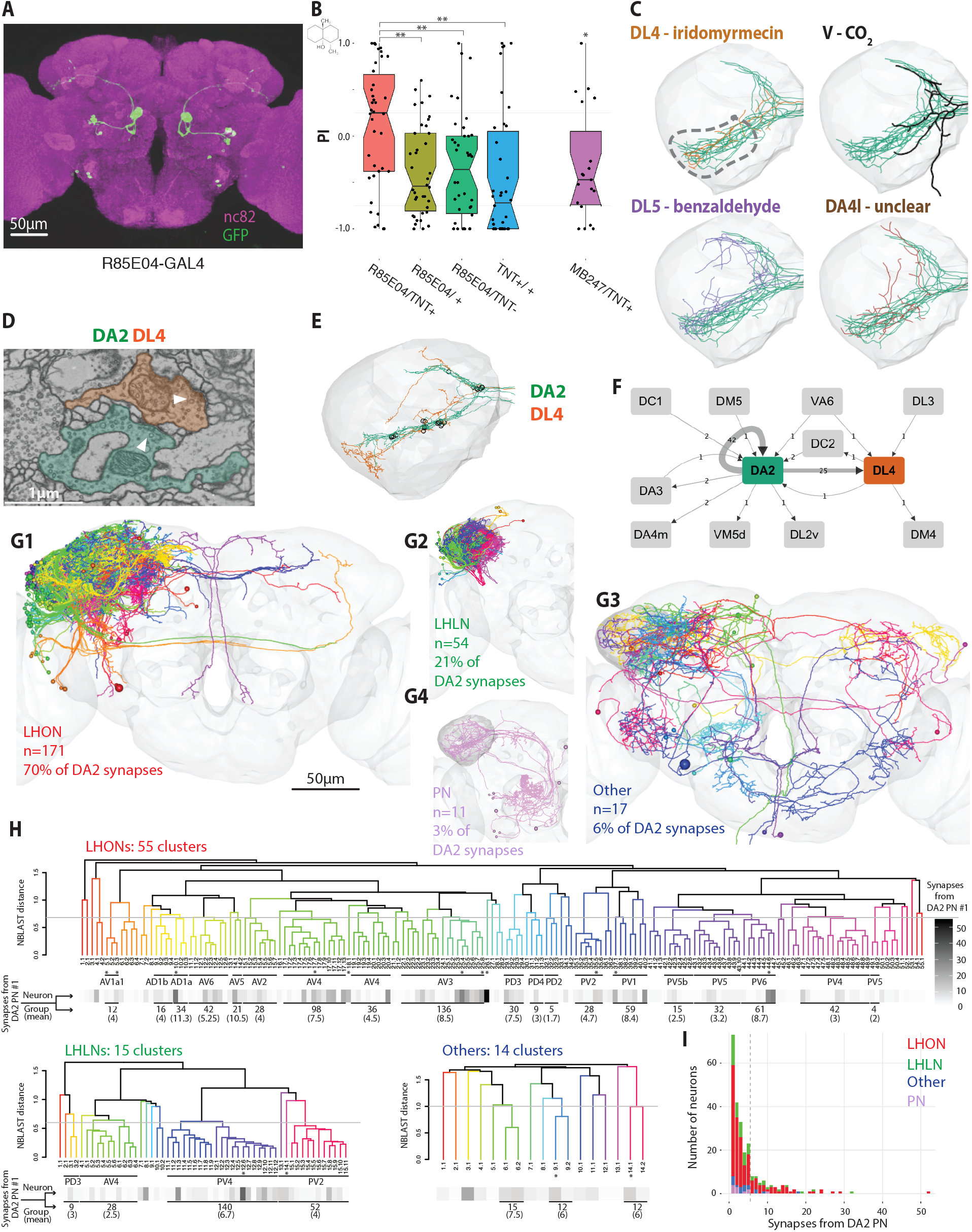
DA2 PNs are necessary for geosmin avoidance, synapse axo-axonically onto the aversive DL4 PNs, and then onto a large number of morphologically diverse third-order neurons. (A) R85E04 labels 5-6 DA2 uniglomerular PNs per hemisphere. (B) Egg-laying two-choice preference indices (PI) to geosmin while silencing DA2 PNs (n=38 for R85E04 and its controls, n=19 for MB247-GAL4/TNT+). (C) Frontal view of DA2 PN axons (green) with examples of other aversive PNs in the LH (DL4, V, DL5). DA4l targets the same area of the LH, but its ligand and valence are unclear. The dashed line marks the putative zone of aversive convergence. (D) An example of a DA2 PN (green) to DL4 PN (orange) axo-axonic synapse in the LH. The arrowheads mark the presynaptic sites. (E) Location of axo-axonic synapses from DA2 to DL4 PNs (open circles). (F) Axo-axonic PN synapses from and to DA2 PNs (n=5) in the right hemisphere. (G1-G4) Downstream targets of DA2 PNs in the LH, by broad neuron class (LHONs, LHLNs, unclassified neurons and PNs). (H) Hierarchical clustering of the DA2 PN down-stream target morphologies. Grey lines mark the cut off heights (LHON:0.68, LHLN:0.60, Others:1.0), greyscale bar under the dendrograms the number of synapses from the completed DA2 PN, and asterisks the completed neurons. (I) Distribution of the synaptic connections from the DA2 PN to its downstream targets, color-coded by broad neuron class. Grey line marks the shoulder of the distribution, between 5 and 6 synapses. Significance values: * p<0.05 ** p<0.01

To demonstrate the functional role of these DA2 PNs, we silenced their synaptic output by expressing tetanus toxin (7) via R85E04. This completely abolished avoidance behaviour (Figure 1B) confirming that DA2 PNs are necessary for geosmin sensing. In summary, R85E04 labels the excitatory DA2 PNs postsynaptic to Or56a ORNs; these PNs respond strongly and selectively to geosmin, and are necessary for geosmin avoidance behaviour during egg-laying, a task of key ethological importance to the animal.

DA2 PNs project to two higher brain areas: the mushroom body (MB) and the lateral horn (LH). The former is thought to be involved in associative learning and the latter in innate behaviour. As expected, the MB appeared not to play a role in geosmin aversion (Figure 1B), so we focused on the LH. The axonal morphology of DA2 PNs shows notable similarities with several other aversive odour processing PNs (Figure 1C), particularly in the ventral-posterior LH (dashed line in Figure 1C), an area recently suggested to be important for egg-laying aversion (8). The similarity is especially striking with DL4 PNs, which are postsynaptic to Or49a/Or85f ORNs tuned to the sex pheromone of the parasitic wasp *L. boulardi* (9). This suggests the possibility of valence-based integration by downstream neurons in the LH (9, 10).

To look for downstream targets of DA2 PNs, and thus possible valence integrator neurons, we took two parallel, complementary approaches. First, we reconstructed DA2 PNs and their postsynaptic partners in the LH in a recently acquired whole-brain electron microscopy (EM) volume of a female *Drosophila* (11), which we refer to as FAFB. We were then able to make comparisons with a second recently acquired partial EM volume of a female (referred to as hemibrain) (12).

Second, we performed a light-level *in silico* anatomical screen to look for genetic driver lines containing LH neurons (LHNs) putatively downstream of DA2 PNs. We will discuss the EM connectomics approach first.

Previous work had already identified all the uniglomerular PNs on the right (R) side of the EM volume (11), including 5 DA2 PNs. We additionally identified 6 DA2 PNs on the left (L) side of the brain, and marked up all their presynaptic sites on both sides (R: 961, L: 959). Surprisingly, we found a relatively large number (R: 25; L: 32) of axo-axonic synapses from the DA2 PNs to the single aversive DL4 PN (Figure 1D, E and F, Figure S2A and B), mostly clustered on the ventral axonal branches (Figure 1E). These axo-axonic synapses are also present in the hemibrain (R: 31) Figure S1G). In FAFB no other PN receives more than 2 synapses from the DA2 PNs (Figure 1F), while in the hemibrain, only one other PN receives more than 4 synapses (DA4m) Figure S1G). In both volumes DA2 PNs also synapse strongly onto each other (Figure 1F, Figure S2B), and Figure S1G), thus showing within-odour channel divergence and re-convergence that may serve to increase signal detection speed (13). As a large number of PNs pass close to DA2 synapses without receiving input (Figure S2C), this connectivity appears specific, not just proximity based. This is also supported by the absence of connections from DL4 PNs to DA2 PNs. Axo-axonic integration of PN odour channels has not been previously described, although our recent work (14) has found that it is widespread within PN axons. This type of connectivity may be an important mechanism for valence-based integration, ultimately triggering similar behavioural responses to odours of similar significance to the fly.

We next obtained a complete downstream connectome of the LH targets for a single DA2 PN (Figure S2D). We reconstructed the postsynaptic partners sufficiently to enable unambiguous identification, and quantification of the DA2 PN input. This identified a surprisingly large array of 253 neurons: 171 LH output neurons (LHONs) (Figure 1G1), 54 LH local neurons (Figure 1G2), 11 PNs (Figure 1G4) and 17 other neurons (Figure 1G3). These last 17 include large brain spanning neurons; 10 appear neuromodulatory due to the presence of dense core vesicles or because they are labelled by the TH-GAL4 driver in the FlyCircuit database (15, 16).

Together, the 225 LHN target neurons make up ~ 18% of the estimated total of ~ 1400 neurons in the LH (17), a very large fraction given that DA2 is just one of 51 olfactory glomeruli (2%). The majority (~ 70%) of DA2 synapses were onto LHONs, and ~ 21% onto LHLNs, roughly matching the proportion of neurons in each broad class. The LHONs project to various neuropils thought to be involved in multimodal sensory integration (Figure S2E) (18–23). The neurons are of diverse morphologies: hierarchical clustering of each broad class, splits them into 84 clusters (55 LHONs, 15 LHLNs, 14 others), excluding PNs (Figure 1H). Taken together this suggests the wiring logic of the LH is massively more complex than previously thought (21, 24): PNs from a ‘labelled line’ glomerulus with clear behavioural meaning do not synapse onto just a few postsynaptic targets. However, the distribution of connectivity is skewed; the majority of targets receive only a few inputs (Figure 1I), and for all 5 RH DA2 PNs in both EM volumes ~ 25% of all DA2 downstream neurons make up more than 50% of all synaptic output(Figure S2D). Moreover, LHONs in the same morphological clusters have a higher than chance probability of getting similar levels of DA2 input (Figure S2F, see Methods), but the same is not true for LHLNs (and was not tested for the other neurons, which are more structurally diverse). In summary, the geosmin processing pathway that starts as a labelled line shows convergence at the level of PN axons with another aversive pathway (DL4 PNs), as well as considerable divergence at the transition from second to third-order level.

In order to answer whether valence-based integration takes place in the LH we selected a sample of 15 DA2 strong downstream neurons of diverse morphologies for complete reconstruction in FAFB. As all the uniglomerular excitatory PNs on the right hemisphere were completed we could identify every PN input onto these neurons. Figure 2A shows the morphologies of the completed neurons (cell typing according to (14)). We observe a range of input tuning profiles, from DA2 specific (AVLP594, a neuropeptidergic brain-spanning neuron, Figure S2G and H (25)) to completely or nearly aversive odour specific (LHPV6a3#1, LHAV3a1_c#1), to relatively broadly tuned (LHAV3f1, LHAV1a1) (Figure 2B). Altogether, 9/15 neurons receive above chance amounts of aversive input from PNs besides DA2. We then identified the same cell types in the hemibrain for comparison. Averaging within cell types and normalising inputs shows few major differences in connectivity for our DA2 downstream targets (Figure 2B). Similar to FAFB, 8/15 neurons in the hemibrain receive above chance amounts of aversive input from non-DA2 PNs.

**Figure 2:**
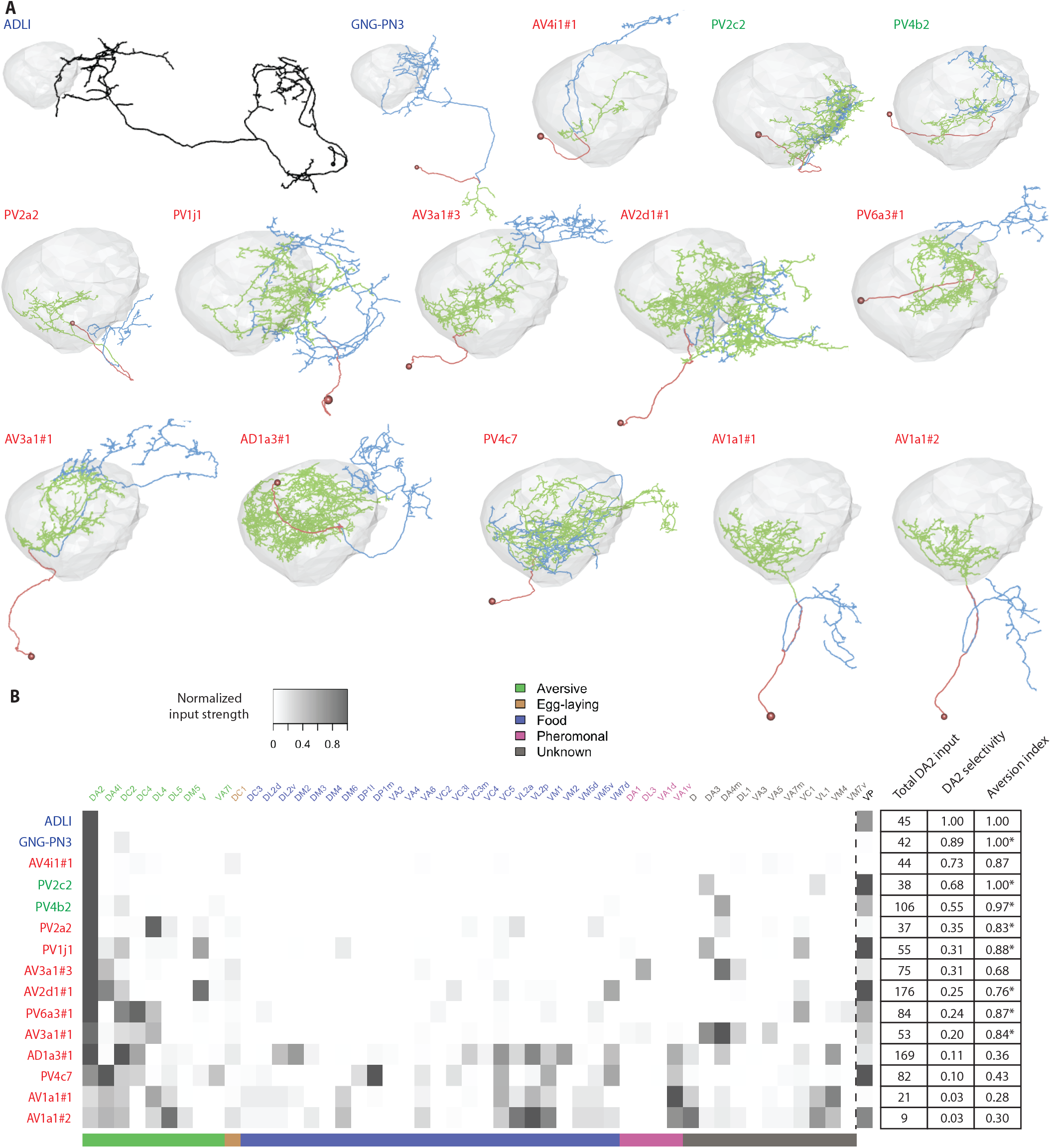
Strong DA2 downstream targets have diverse tuning breadths but tend to receive more than a chance amount of non-DA2 aversive PN input. (A) EM-reconstructed morphologies of selected strong DA2 targets. Cell bodies and primary neurites are coloured in red, dendrites in green, and axons in blue, where polarity is clear. Neuron names are color-coded by broad neuron class (other: dark blue, LHON: red, LHLN: green). (B) A heatmap representation of excitatory uPN to LHN connectivity for top DA2 targets in both EM volumes, FAFB and the hemibrain, normalised by the total uPN inputs to each neuron, with glomeruli color-coded by putative behavioural relevance ((2, 9, 27–35)). The total amount of DA2 input, DA2 selectivity (DA2 input/total uPN input), and Aversion index (DA2 input/uPN input from PN channels with known valence). Neurons receiving more than chance amount of non-DA2 aversive input are marked with an asterisk.

In addition, most of these neurons receive input from thermo- and hygrosensory VP glomeruli (Figure 2B) (26). This may reflect either direct multisensory integration of aversive signals for extremes of temperature or humidity (critical dangers for insects), or cross-sensory modulation of olfactory pathways by environmental context. These patterns of synaptic connectivity therefore support valence-based integration occurring in the LH. However this is unlikely to be the only computation taking place at this transition from second to third-order level of the circuit.

In parallel with our EM work, we carried out a light-level screen for DA2 downstream neurons and driver lines. We used a registered confocal stack of R85E04 and converted the LH axon arbour into a binary mask (Figure 3A). This allowed us to identify sparse driver lines from the GMR-GAL4 (5) and LH-Split collections (36) containing LHNs with dendrites overlapping the DA2 axons (Figure 3B and C). This *in silico* anatomical screen identified 18 LHN types, 12 of which could be accessed relatively specifically through either GAL4 or Split-GAL4 lines.

**Figure 3:**
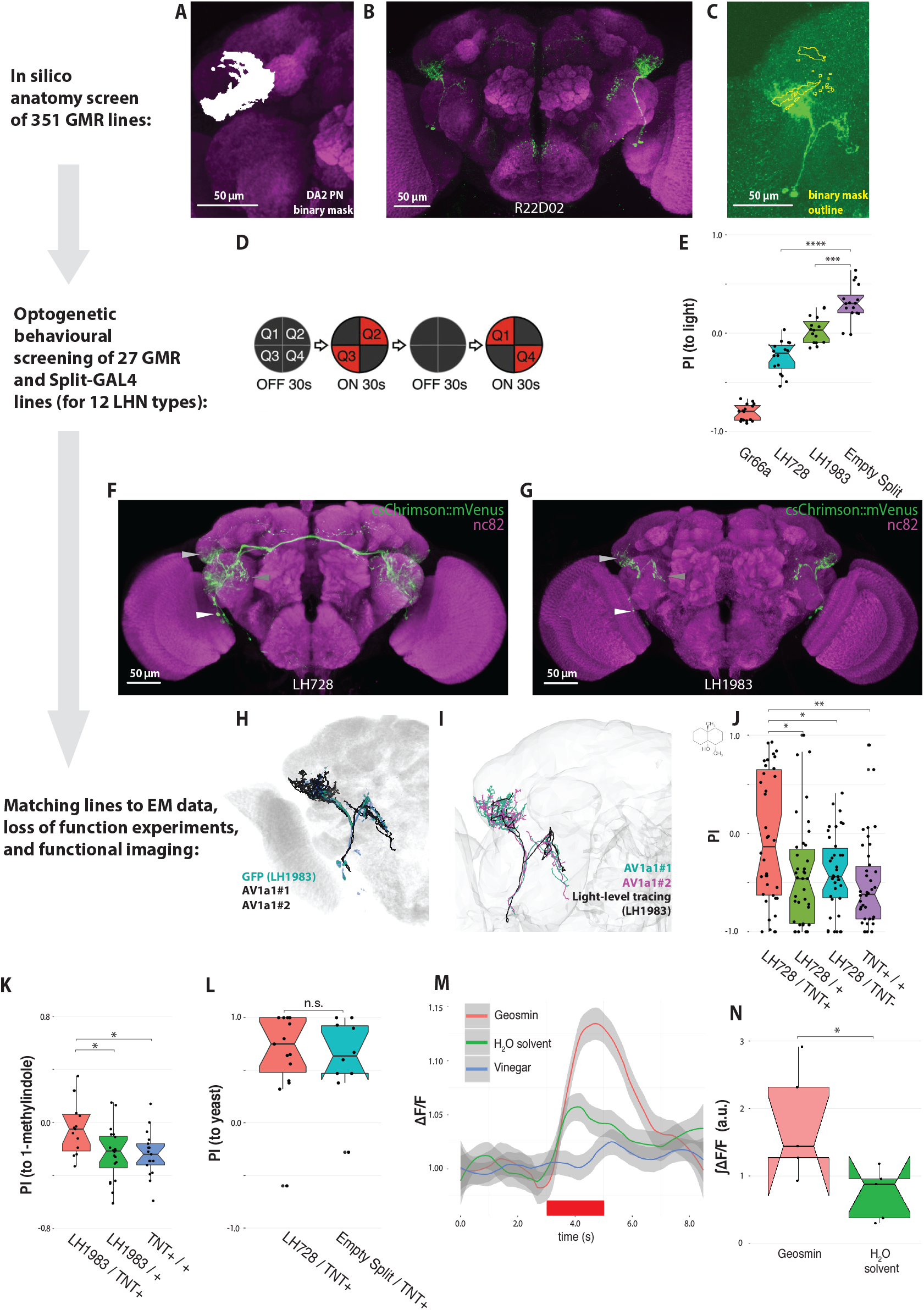
LHAV1a1 neurons as sufficient for aversion and necessary for both geosmin and 1-methylindole avoidance. (A) LH close-up of the binary mask of DA2 PNs. (B) An example of a good hit in the *in silico* screen. (C) An example of the binary mask (as an ROI, in yellow) together with a good hit. (D) A schematic representation of the optogenetic four quadrant assay ((37)) used for behavioural screening. 20 female flies explore a circular arena for 30 seconds before two of the quadrants are illuminated with red light for 30 seconds, after which the protocol is repeated by illuminating the remaining two quadrants. Reproduced with permission from ((37)). (E) PI (to the light quadrants) for the last 5 seconds of the stimulation epochs (n=16, for all groups). (F-G) Expression patterns of LH728 and LH1983. The arrowheads mark the cell bodies (white), dendrites (light grey), and axons (dark grey) of the type LHAV1a1 LHNs. (H) A 3D rendering of EM reconstructions of LHAV1a1#1 and #2 (black) overlaid with the LH1983 expression pattern (green). (I) A light-level tracing of the LHAV1a1 LHNs (black) overlaid with EM-level tracings of LHAV1a1#1 and #2 (green and magenta, respectively). (J) Egg-laying two-choice PI to geosmin while silencing LHAV1a1 LHNs (n=38 for all groups). (K) Egg-laying two-choice PI to 1-methylindole in the split-plate assay while silencing LHAV1a1 LHNs (n=15-20). (L) Egg-laying two-choice PI to yeast odour while silencing LHAV1a1 LHNs (n=14 for both). (M) Odour-evoked *in vivo* two-photon imaged calcium responses (GCaMP3.0) from LH dendrites of LHAV1a1 neurons (n=5). Red bar marks the stimulation epoch. (N) Area under curve for the stimulation epoch for geosmin and its solvent control (green). Data same as M (n=5). Significance values: * p<0.05 ** p<0.01 *** p<0.001 **** p<0.0001

With these reagents in hand, we carried out an optogenetic activation screen hoping to identify aversion triggering LHNs. A total of 27 driver lines (for 12 LHN types) were tested using a four-field arena (Figure 3D, see (36) for full details of the screen and (37) for the apparatus). Only two of the tested driver lines triggered aversion in this assay (Figure 3E and Figure S3A, B and C): LH728 and LH1983 (Figure 3F and G). Both lines share a parental line (R76E07) and label ~ 10 neurons, with somas ventral and medial to the anterior ventrolateral protocerebrum (AVLP). Interestingly, both lines also contain the same two neurons with dendrites in the (ventral) LH and axons in the AVLP, and no other LHNs. The neurons are cholinergic (36), and appear morphologically very similar to the LHAV1a1 neurons found downstream of DA2 PNs in the EM volume (Figure 3H). To confirm this, we generated light-level tracings of the neurons in LH1983. As the processes of the two LHNs were in many places too close to resolve, the tracing resulted in a single hybrid skeleton of the two AV1 neurons found in the line. However, overlaying the light-level tracing with LHAV1a1#1 and LHAV1a1#2 reveals a remarkably similar morphology (Figure 3I). Moreover, a quantitative NBLAST (16) comparison to all the 33 neurons taking the AV1 tract in the EM volume shows that the tops matches are LHAV1a1#1 (similarity score=0.69) and LHAV1a1#2 (similarity score=0.65), respectively. Intriguingly, These cells are one of only 3 out of 70 LHN cell types downstream of DA2 PNs that we identified with projections to the ventral rather than superior protocerebrum in the FAFB volume. There are three groups of AV1a-like neurons in the hemibrain, belonging to three classes: AVL02q_a_pct (3 cells), AVL02q_b_pct (4 cells) and AVL02q_c_pct (2 cells). A comparison of morphology and olfactory inputs shows that AVL02q_c_pct are the most similar to FAFB LHAV1a1s (Figure 3I and J).

We also tested whether the LHAV1a1 neurons are necessary for geosmin avoidance by silencing their synaptic activity using tetanus toxin; this abolished geosmin avoidance in the egg-laying assay (Figure 3J). *In vivo* calcium imaging confirmed that the neurons respond to geosmin, but not to vinegar (a broadly coded attractive odorant) (Figure 3M and N). Taking these functional data together with the fact that no other AV1 tract neurons receive a significant amount of DA2 PN synaptic input in EM data, strongly suggests that the LHAV1a1 LHNs labelled by both driver lines are necessary and sufficient for some forms of odour avoidance.

Are the other 69 classes of lateral horn neurons that receive geosmin information then irrelevant to its processing? There are a number of possible explanations that could explain this large number of circuit elements.

The input tuning of the pair of LHAV1a1 neurons that we reconstructed is relatively broad, receiving input from multiple aversive PN channels (including DA2 and DL4). We therefore wondered if these neurons have a general role in aversive odour processing. There are relatively few low concentration repulsive odorants reported in *Drosophila*. The DL4 ligand iridomyrmecin (9) is not commercially available and pilot control tests with a limited amount of synthetic compound (kindly shared by Jerrit Weißfog and Ales Svatos) did not show robust aversion in our egg laying assay (data not shown). We therefore tested another compound, 1-methylindole, a derivative of the bacterial metabolite indole (38), which we found to be aversive. Silencing LHAV1a1 neurons also abolished this egg-laying aversion (Figure 3K, see also Figure S3D-F, and methods). Indoles are detected by a phylogenetically closely related receptor Or43a (39, 40), which projects to the DA4l glomerulus (40). Intriguingly LHAV1a1 LHONs also receive strong input from DA4l PNs (Figure 2B). Importantly, flies were still attracted to yeast odour (Figure 3L), showing they are not anosmic or otherwise unable to respond to odours. Together these data provide evidence for functional integration of aversive odour channels within LHAV1a1 neurons, which appear to have a selective role in odour avoidance during egg-laying.

Combining functional and connectomics approaches, provides an almost unique opportunity to explore the brainwide logic of olfactory processing. However, the considerable divergence of the geosmin processing pathway when moving from second to third-order neurons, means that it is presently impossible to follow all of the connected LHNs deeper into the brain. Instead we focussed on two cell types that appeared of special interest from the results so far: LHAV1a1 and PV6a3 (an ideal example of an aversive integrator based on its PN inputs).

We identified 14 postsynaptic partners receiving 2 or more synapses downstream of PV6a3, the strongest of which appears to be a neuron projecting to the suboesophageal zone (SEZ) (Fig S4A, B and C). However, as we did not perform an exhaustive reconstruction of the postsynaptic partners, there most likely are more than shown here. Nevertheless, this suggests that one next computational step in the circuit is to integrate aversive odour signals with gustatory ones in the SEZ. Downstream sampling from the LHAV1a1#1 axon identified 44 postsynaptic partners, many shared with its sibling LHAV1a1#2 (Figure 4A, Figure S4D and E). Downstream sampling in both connectomes showed that LHAV1a1s share many postsynaptic partners, and, most notably, are all strongly connected to descending neurons (DNs)Figure S4C and D), including two previously unreported DNs, which we have named DNp42 and DNp43, projecting to the nerve cord (Figure 4A). LHAV1a1 form many more direct connections with DNs that our other strong DA2 targets Figure S4. Another strong target is DNp06. However, while it receives ~ 5.4% of LHAV1a1 output, LHAV1a1 input to this DN comprises just 0.3% of the total.

**Figure 4:**
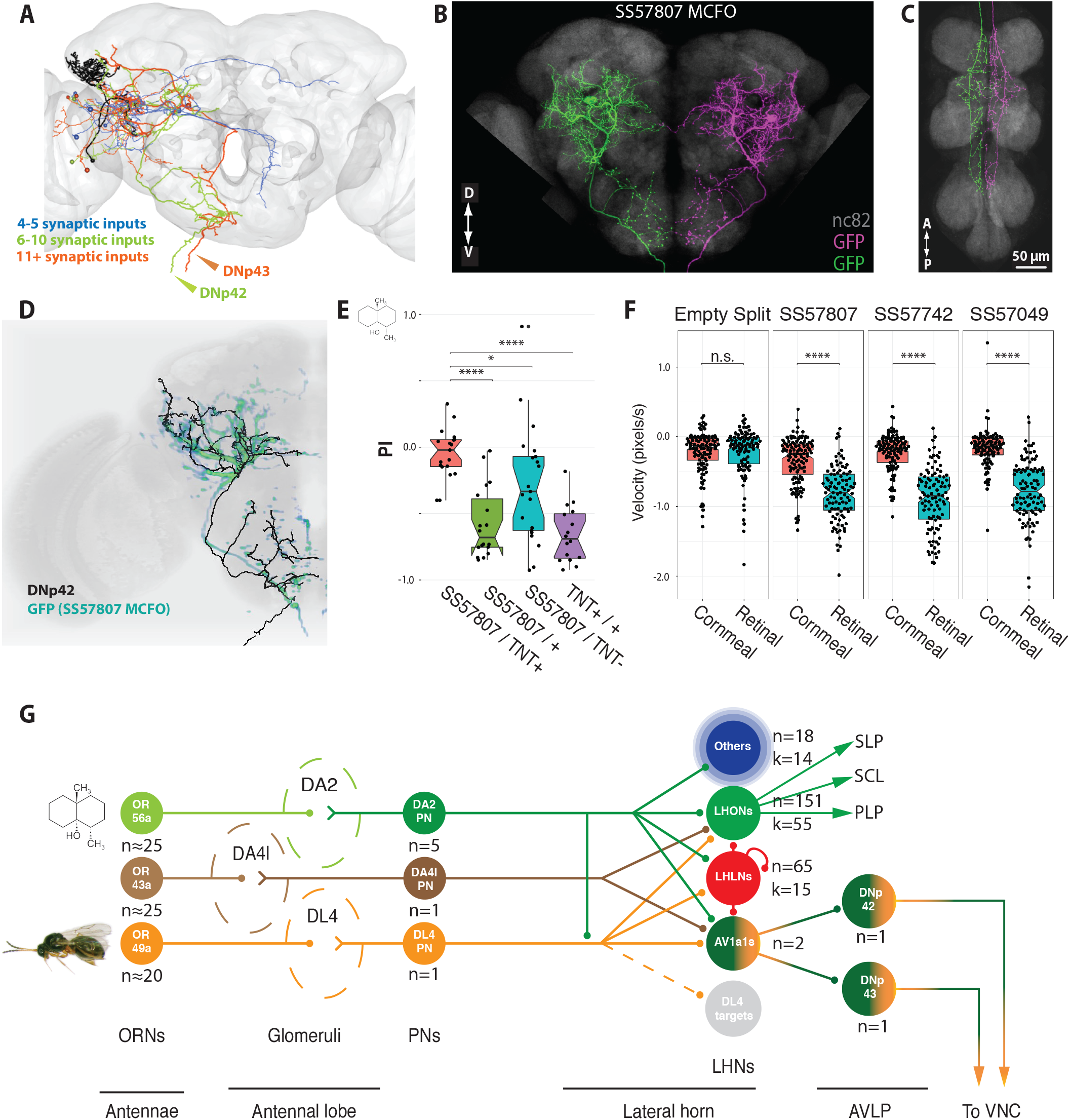
LHAV1a1 neurons synapse onto DNs that are necessary for geosmin avoidance and sufficient to trigger avoidance behavior. (A) Identified downstream targets of the LHAV1a1#1 with 4 or more synaptic inputs. Neurons are colour-coded according to number of synaptic inputs. Two descending neurons DNp43 (orange) and DNp42 (green) are highlighted with arrowheads. (B-C) MultiColor FlpOut (MCFO) labelling of putative DNp42 neurons in the brain (B) and VNC (C). (D) A 3D rendering of EM reconstruction of DNp42 (black) overlaid with the SS57807 expression pattern (green). (E) Egg-laying two-choice PI to geosmin while silencing DNp42 neurons (n=16-21). Fly velocity in response to optogenetic stimulation of DNp42 neurons via three different driver lines (n=99-109). Significance values: * p<0.05 **** p<0.0001 (G) A simplified schematic of the circuit. Geosmin is detected by Or56a ORNs that project to the AL where they synapse onto DA2 PNs, and local neurons (not shown). The DA2 PNs project to the LH (forming en passant synapses in the MB, not shown). In the LH, DA2 PNs form axo-axonic synapses with parasitic wasp pheromone processing DL4 PNs, and then synapse onto a large number of third-order neurons with varying tuning breadths. LHAV1a1 neurons receive input from multiple aversive PN channels (including DL4 and DA4l), and synapse onto DNs that trigger avoidance. Numbers of neurons (n) and clusters (k) are marked.

We identified three sparse driver lines for DNp42 (Figure 4B-D, and Figure S5A-C). Silencing DNp42 abolishes geosmin avoidance (Figure 4E), reproducing the phenotype seen at every step in this circuit, from sensory input to these descending neurons. We also optogenetically activated these DNs while observing the flies using high-speed videography (FlyPEZ assay (41)). Strikingly, light-activation triggered a consistent backing up phenotype (Figure 4F) and flies would occasionally take-off, similar to what is seen in response to looming visual stimuli (42). In the VNC, DNp42 axons arborise in all three thoracic neuromeres but stay close to the midline innervating the tectulum (an integrative area), avoiding the more lateral leg neuropils. They also innervate the accessory mesothoracic neuromere and ventral association centre, areas receiving sensory input from the wings and legs, respectively (43) (Figure S6A and B). There are no projections to the abdominal ganglion (which directly controls reproductive functions, including egg laying) and no obvious sexual dimorphism in any of these axonal arbours (Figure S6C). This anatomy is consistent with a pre-motor and/or sensory-motor integration function in locomotor behaviour rather than a direct impact on motoneurons or regulation of egg laying.

While in FAFB there is no exhaustive upstream tracing of DNp42, the autosegmentation of the hemibrain allows us to see it’s other inputs. LHAV1a1s comprise ~ 1.3% of all input to DNp42. The second strongest input to DNp42, comprising ~ 2.6% of all input, is a GABAergic mushroom body output neuron, MBON20(y1y2). This cell arborises in the y1 and y2 lobes of the ventral accessory calyx of the mushroom body. These lobes receive axonal input from protocerebral posterior lateral dopaminergic neurons, which carry predominantly aversive signals. This connectivity resembles the circuit for learned-innate interaction described in (44), though at the level of DNs, rather than LHNs, and with the opposite valence, possibly providing a mechanism by which learned and innate olfactory information is integrated to produce appropriate behavioural responses.

All animals need to solve similar challenges to survive and reproduce in a complex environment. Avoiding pathogens and parasites are some of the ubiquitous ones. Here we trace a microbial-odour processing pathway from sensory neurons all the way through the brain to descending neurons innervating the fly’s homologue of the spinal cord (see Figure 4G, for a summary of the circuit). We have identified circuit motifs including multiple levels of valence specific convergence, including clear evidence for convergence of aversive PNs onto common targets, which tend to be co-located in space. This supports the idea of valence-based topography as one organising principle of the LH (8, 10, 45). More surprisingly, we also found that central olfactory layers show a highly diverging organisation, even for stimuli that are initially coded in a labelled line fashion. This contrasts with our functional data: we trace a single linear pathway essential for aversive egg-laying behaviour, as supported by neuronal activation and/or silencing experiments at each of the 4 layers traversing the brain. Although our analysis of fourth order neurons is far from exhaustive, we have identified few other direct connections to a descending neuron. Therefore this synaptic pathway is probably unusually shallow. This in turn may imply both a particular biological significance and make these neurons more sensitive to simple experimental activation or silencing experiments.

Comparing our connectomics and experimental results challenges us to think about what connection strengths are behaviorally relevant. (46) has recently suggested a classification of major pathways, with 100 or more inputs, and minor pathways, with fewer than about 10. Summing inputs from all 5 DA2 PNs we identified just three target neurons meeting this criterion (range 107-176 connections), not including the LHAV1a1s, which are individually only the 42nd and 90th strongest DA2 targets by synapse number. Normalising by the total number of inputs, in the majority of cases DA2 PNs accounted for <5% of the input to target neurons, and just 1.6% of the input to the reconstructed LHAV1a1 neurons. Thus, while absolute thresholds are a useful rule of thumb, it is likely that different criteria will be necessary in different brain areas and between different kinds of neurons. This will be of major significance in interpreting the dense whole brain connectomics datasets in *Drosophila* recently released (12), with larger brains likely following within the space of a few years (47).

## Materials and Methods

### Drosophila stocks, driver line generation, and husbandry

The following stocks were used: Canton S (UC San Diego *Drosophila* Stock Center, CA); Ir8a1; Ir25a2; Orco1, Gr63a1 (a kind gift from R. Benton); w; Or56a-GAL4; + (Bloomington *Drosophila* Stock Center, Indiana); w; Or56a-/-; + (a kind gift from Christopher Potter) (8); w; MB247-GAL4; + (Bloomington *Drosophila* Stock Center, Indiana); w; UAS-Kir2.1::GFP;+ (a kind gift from Matthias Landgraf); w; +; R85E04, Bloomington *Drosophila* Stock Center, Indiana) (5); w; UAS-TNT-active form; w; UAS-TNT-inactive form (both kind gifts from C.O’Kane) (7); 20XUAS-IVS-ChrimsonR::mVenus (attP18);+;+ (48); w; UAS-GCaMP3.0 (attP18); UAS-GCaMP3.0 (attP40); w; UAS-mCD8::GFP; UAS-mCD8::GFP; +; y, w; poxnMB00113; + (Bloomington *Drosophila* Stock Center, Indiana); *teashirt*-GAL80 (49); Zp-GAL4DBD, pJFRC200-10XUASIVS-myr::smGFP-HA (attP18); +; +; p65ADZp(su(Hw) attP8); +; +; w;p65ADZp(su(Hw) attP40); +; w; +; p65ADZp(su(Hw) VK00027); pJFRC51-3xUAS-Syt::smGFP-HA ((Hw) attP1); pJFRC225-5xUAS-IVS-myr::smGFP-FLAG (VK00005).

The LH Split-GAL4 (50) lines for the optogenetic screen were made as a part of a larger collaborative screen for creating a cell-type specific driver line library for LHNs (details described in (36)), using a subset of the enhancer fragments used in generating the original GMR GAL4 lines (5). The Split-GAL4 lines for the DNs were made essentially similarly to (51). Based on our screening of GAL4 and GAL4 with *teashirt* lines, we selected AD/DBD combinations from the Janelia (52) and VT (53) collections that we thought shared expression in individual DNs. To visualize combined expression patterns, we crossed males carrying a GFP reporter (pJFRC200-10XUASIVS-myr::smGFP-HA in attP18) and the ZpGAL4DBD transgene (in attP2) with virgin females carrying the p65ADZp transgene in either su(Hw)attP8, attP40, or VK00027 and examined expression in 3- to 10-day-old female progeny. The split-GAL4 combinations that we deemed sparse enough to include in our DN collection were made into stable stocks containing the AD and DBD transgenes. To obtain polarity and higher resolution (40x, 63x) information on selected lines, split-GAL4 lines were crossed to pJFRC51-3xUAS-Syt::smGFP-HA in su(Hw)attP1; pJFRC225-5xUAS-IVS-myr::smGFP-FLAG in VK00005 and processed for imaging. We used the multicolor flip out technique to stochastically label individual neurons in lines that contained multiple cells (54). These protocols are available on the Janelia FlyLight website (https://www.janelia.org/project-team/flylight/protocols). Some split-GAL4 lines were also crossed to 20XUAS-CsChrimson-mVenus trafficked in attP18 (virginator stock) and processed as above to visualize expression pattern when using the CsChrimson effector, as observed expression patterns are known to vary slightly depending on the reporter used (55). Based on their GFP or CsChrimson expression patterns, we made our best estimate of the number of background (non-targeted-DN) cell types in each split-GAL4 line made, and we gave each split line a quality score of A (no background expression), B (one background cell type), or C (two or more background cell types). Confocal image stacks of the stabilized split-GAL4 intersections are available online (http://www.janelia.org/split-gal4). For most experiments flies were reared at 25 C and 60% humidity, under a 12:12 hour light-dark cycle, on food made with the following recipe: 4.8 l H2O, 275 g of Glucose, 250 g yeast, 37 g agar, 175 g flour, 125 ml Nipagen solution, 50 ml penicillin/streptomycin, 20 ml propionic acid. The same food was also used for the egg-laying assays. For optogenetic behavioural experiments, flies were reared at 22C on standard Iberian food containing yeast, cornmeal and agar, and supplemented with 1/500 all-trans retinal (Sigma-Aldrich, St. Louis, USA).

### Odor stimuli

Geosmin (CAS #16423-19-1) was used at a concentration of 1:1000 (Sigma-Aldrich, St. Louis, USA, Product Number UC18). 1-Methylindole (CAS #603-76-9) was used at either 1:1000 or 1:10.000 concentration (Sigma-Aldrich, St. Louis, USA, Product number 193984). The odors, concentrations, and odor delivery used for electrophysiology and calcium imaging were the same as used in (17).

### Egg-laying two-choice assay

Female flies were collected on the day of eclosion under CO2 anaesthesia, reared in same sex vials at 25 C and 60% humidity. Female flies aged 5-7 days were then mated with males of similar age for 6 hours on the day of the experiments. After six hours of mating the female flies were again isolated from the males under CO2 anaesthesia and were left to recover for 2 hours before starting the experiments. For the experiments, approximately 20 females were transferred without anaesthesia into a BugDorm insect rearing cage (24.5×24.5×24.5 cm) (MegaView Science Co., Ltd., TAIWAN) made of polyester netting. Two ø 50mm Petri dish plates containing Iberian fly food were placed in opposing corners of the cage and a small plastic cup cut from the cap of a 1.5 ml Eppendorf tube containing the experimental odour, or the solvent control, was placed at the center of each food plate. For the geosmin experiments 5 μl of geosmin (1:1000 dilution in mineral oil) was used as a stimulus. For the experiments done with yeast odour, 100 μl of 400 mg/ml of baker’s yeast in Milli-Q H2O was used. A nylon mesh was used to physically separate the flies from the odorant. All experiments started at 12:00 h Zeitgeber time (+/− 1 h) and lasted for 16 hours (+/− 1 h). Eggs were counted under a stereo microscope. An oviposition Preference Index (PI), was calculated by using the formula

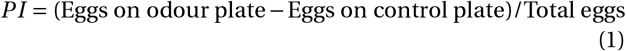

PI could thus get values from −1 to +1, signifying total avoidance and total preference of the geosmin plate, respectively.

We also developed an alternative version of the egg-laying assay (adapted from (31) and used in Figure S3D, E and F, Figure 3K, and Figure 4E). For this we used a single ø 50mm Petri dish plate containing Iberian fly food. The plate was split in two by a divider, the food was then melted by briefly heating the plates, and the experimental odour (50 μl of 1:1000 geosmin in Milli-Q H2O) and solvent control were mixed directly into the food. After the food had solidified again, the divider was removed. Five mated females were aspirated onto the plates and the Petri dish plate was placed back on top of the plates. Experimental duration and other parameters were as above. The main benefit of this version of the assay was that the PI variance was lower, which allowed us to use i) lower sample sizes and, ii) fewer flies per replicate, thus leading to a significantly improved experimental throughput. We ascertained that the behavioral phenotype was still solely due to olfaction (Figure S3D), and replicated the main results we obtained with the other assay (Figure S3E). PI was calculated as

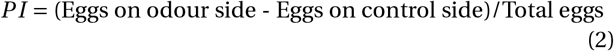

For the 1-methylindole experiments, we tried both 1:1000 and 1:10.000 concentrations. As a two-way ANOVA with genotype and concentration as factors showed a significant main effect for genotype, but not concentration, and there was no observable genotype × concentration interaction, we pooled both concentration groups for Figure 3K, with ~ half of the flies for each genotype coming from each stimulus concentration.

### Egg-laying no-choice assay

Fly collection, rearing and mating was performed similarly to the two-choice assay. For the no-choice assay, 5 females were aspirated without anaesthesia onto ø 50 mm Petri dish plates containing fly food, and the lid was placed on the plate. In the experiments where the effect of odorants on egg-laying quantity was tested, the stimuli were pipetted onto a small plastic cup cut from the cap of a 1.5 ml Eppendorf tube. 50 μl of geosmin was used as a stimulus. A nylon mesh was used for physically separating the flies from the odorant. All experiments started at 12:00 h Zeitgeber time (+/− 1 h) and lasted for 16 hours (+/− 1 h). Eggs were counted under a stereo microscope.

### Optogenetic four-field assay

The four-field optogenetic assay was carried out essentially as described in (37). Crosses were made on normal fly food containing 1:500 all-trans retinal (Sigma-Aldrich, MO, USA), and eclosed females of the right genotype were collected into same-sex vials under cold anaesthesia, and reared in the dark on 1:250 all-trans retinal food, at 22 C and 50% humidity. Approximately 20 female flies, aged 3-7 days were used for each experiment. The females were not specifically mated for the experiments, but they were producing fertilised eggs by the time of the experiments. The assay was performed on a circular arena of 10 cm diameter, and 3 mm height. Flies were transferred onto the arena without anaesthesia by using a vacuum pump. All experiments took place in darkness, at 25 C and 50% humidity. To prevent the infrared backlight from affecting the temperature in the arena, the arena was mounted on a heat sink, and an airflow of 150 ml/min from the four corners of the arena to the centre was maintained.

A 617 nm wavelength LED (Red-Orange LUXEON Rebel LED; Luxeon Star LEDs, Brantford, Ontario, Canada) was used for the optical stimulation. The behaviour of the flies was recorded by using a camera (ROHS 1.3 MP B&W Flea3, US 3.0 Camera; Point Grey, Richmond, BC, Canada) equipped with a long-pass (800 nm) filter (B&W filter; Schneider Optics) set to capture at 30 Hz, and controlled via a custom script written in Matlab. Only water was used for cleaning the arena.

### FlyPEZ assay

The FlyPEZ assay was carried out as described in (41).

### Neuronal reconstructions in EM

Two EM datasets were used: a full female adult *Drosophila* brain (FAFB) (11), and another partial adult female brain (hemibrain) (12). In FAFB, neuron skeletons were manually traced using CATMAID (56, 57), following the procedure as described in (11). The identification of antennal lobe glomeruli and their cognate PNs follows (14). PNs were traced to completion in the LH, and all their presynapses and postsynapses were annotated. In the hemibrain, initial neuronal morphologies were generated by machine learning methods and then proofread by human experts. This process resolved any mistakes and merged additional processes, although neurons were not finished to completion. The average neuron completion rate in the LH, as measured by the percentage of postsynaptic densities that belong to morphologies of a significant size, is only 19%. The identification of synapses was an entirely automatic process (12) that differs from the synapse annotation process in CATMAID. For the reasons above, comparing connectivity between FAFB the hemibrain as to be done with care. The identification of PNs and LHAV1a1 neurons is as described in Scheffer *et al.* (12). The neuron morphologies and connectivity used are as released in [neuPrintExplorer version 1.1](https://neuprint.janelia.org/)

### DA2 PN downstream sampling in FAFB

DA2 PN synapses were identified, and all postsynaptic nodes were annotated (consistent with the criteria described in (11)). Once completed, the full set of postsynaptic nodes for a single representative DA2 PN was randomised. Each post-synaptic node was then used as a starting point for tracing out a downstream partner. This sampling procedure was continued until all postsynaptic nodes were either connected to identifiable neurons, or excluded from further analysis due to not being able to connect it to a neuronal backbone (defined by the presence of visible microtubules) as a result of ambiguous features or missing EM sections. Overall, 74.11% of postsynaptic nodes were connected to an identifiable neuron.

Partner neurons were initially traced just enough to identify a soma, thus confirming whether the starting node belonged to a new or previously traced neuron. A sample of 15 LHNs of particular interest (the set in Figure 2) were then traced ‘to completion’; all identifiable branches of the neuron were fully traced, and all incoming and outgoing synapses annotated (note: the neuropeptidergic brain-spanning neuron AVLP594 was only fully traced in the LH). With tracing completed, we were able to examine these neurons’ complete morphology (manually identifying the primary neurite, dendrites, and axon for each), as well as their PN inputs within the LH.

This basic sampling procedure was repeated for the axons of selected third order neurons (LHAV1a1#1 and PV6a3#1). However, the process was not continued to completion for these neurons (Figure S4A and D).

### Potential DA2 axo-axonic connectivity in FAFB

To assess whether or not the DA2-DL4 connectivity was specific, we checked which PN skeletons pass within 1μm of a DA2 output synapse in the LH (in the right LH, approximately 86% of PN-PN axo-axonic synapses occur within this thresh-old—data not shown). Each instance where a PN skeleton was within this 1μm radius was counted as a single potential synapse. This potential connectivity was then compared to the observed DA2 PNs axo-axonic connectivity.

### Clustering of FAFB neuron morphologies

Neurons downstream of the completed DA2 PN were first divided into four broad groups: LHONs, LHLNs, PNs, and others. PNs were excluded from further analysis, and each broad group of neurons analysed separately. The nat.nblast package (https://github.com/natverse/nat.nblast) was used for both NBLASTing (16, 58, 59) and hierarchical clustering of the neurons. More specifically, within the broad groups, each neuron was split into a primary neurite (approximated by taking the longest unbranching segment of the neuron), and the rest (the complement of the primary neurite approximation). Both parts of each neuron were then converted to dot properties representations (16, 58). The NBLAST algorithm (16, 58) was used to generate two all by all similarity matrices; one for the primary neurites, and one for the rest of the neurons. The obtained matrices were then combined by taking the weighted element-wise mean of both matrices, so that the primary neurite was assigned a weight of 0.8 and the rest of the neuron a weight of 0.2. This was done to i) more closely match the manual annotation system of LHNs (that uses the primary neurite tract as the first distinguishing feature in a tri-level hierarchical scheme) (17), and ii) deal with the incompleteness of a large proportion of the neurons. The NBLAST similarity matrices were then converted to distance matrices, and hierarchical clustering was performed by using the average linkage method. Cut heights were determined separately for each broad group (LHON, LHLN, other) after manually assessing the cluster groups.

### Neuron and neuropil nomenclature

Annotation of neuronal types was based on Bates *et al.* (14) and Scheffer *et al.* (12)(for LHNs) and Namiki *et al.* (51) (for DNs). For the cases in which there is more than one individual per type, each individual has been given a unique name by adding ‘#<number>’ after the type name. Image data for light level type example skeletons from FlyCircuit for previously described types (15) can be browsed by searching for the neuron identifier at http://www.virtualflybrain.org/. Neuropil nomenclature was based on (18) for the brain, and (60) for the VNC.

### *In silico* anatomy screen

A confocal stack of an R85E04 brain was registered onto a JFRC2 template brain (5), and the axonal arbors of the DA2 PNs were converted into a binary mask with the Segmentation Editor tool in the Fiji software (NIH, Bethesda, USA). The mask was then used as an ROI to look for driver lines with expression overlapping with the DA2 axons in the LH. This was done by overlaying Janelia FlyLight GAL4 lines (5) with expression in the LH with the mask and visually assessing the overlap. Each line was scored for goodness of overlap, and the neurons in the best lines were identified (17), and then cross-identified in the LH Split-GAL4 lines (36), where possible.

### Confocal microscopy

A Zeiss 710 confocal microscope was used for image acquisition. Brains were imaged at 768 × 768, or 2048×1024 (AL closeup), pixel resolution in 1 μm slices (voxel size: (0.46 × 0.46 × 1 μm) using an EC Plan-Neofluar 40x/1.30 oil immersion objective (Carl Zeiss AG, Jena, Germany) and 0.6 zoom factor. All images were acquired at 16 bit colour depth.

### Image registration

Image registration for the confocal data was carried out according to (61). In brief: the presynaptic marker Bruch-pilot (labeled by nc82) was used as a basis for performing an intensity-based non-rigid warping registration (62) onto a template brain (JFRC2 or JFRC2013, available here: https://github.com/jefferislab/BridgingRegistrations).

The registration procedure itself was performed by using the cross platform Computational Morphometry Toolkit software (http://www.nitrc.org/projects/cmtk). Bridging registrations were used for transforming neurons from one template brain to another (11, 59) by using the nat.flybrains (https://github.com/natverse/nat.flybrains) and elmr (https://github.com/natverse/elmr) R packages. A similar template was derived from the nc82 expression pattern in the VNC of an example female Canton S fly imaged by the FlyLight Project team (template available here: https://github.com/VirtualFlyBrain/DrosAdultVNSdomains/blob/master/template/Neuropil_185.nrrd). Our VNC alignment pipeline was adapted from (60). Briefly: confocal VNC stacks were first converted to an 8-bit nrrd file format, preprocessed using the nc82 reference channel to normalize contrast across samples, rotated to approximately orient the VNC along the anterior-posterior axis, and then the channels were aligned to the template by nonrigid warping (62) using the Computational Morphometry Toolkit.

### Image processing for DN lines

Neuron tracing was carried out semi-manually using Amira 5.4.3 (Visage Imaging, Fuerth, Germany). Volume rendering was performed using Amira ‘generate surface’ function. We first detected the signal with the Amira ‘Interactive Thresh-olding’ function. We then corrected any false detection by manual tracing. Using this image as a mask, we obtained the final masked images shown in the figures using a custom-made program written in MATLAB and the image processing toolbox (MathWorks, Natick, MA, USA). The contrast and brightness of images were modified in Image J (National Institutes of Health, Bethesda, MD). Confocal image stacks of split-GAL4 expression patterns in the brain were aligned to standardized brain template JFRC2013 (see above).

### Light versus EM comparisons of neuron morphology

All neurons taking the AV1 tract in EM were traced far enough to identify major branches and overall morphology. The light-level tracing of LHAV1a1 was obtained by semiautomated tracing in Amira (Thermo Fisher Scientific) from LH1983. Both light and EM neurons were converted to the FCWB reference space via nat.flybrains (https://github.com/natverse/nat.flybrains) (59), and elmr (https://github.com/natverse/elmr) packages for R, and then to dot properties representations by the nat.nblast package (https://github.com/natverse/nat.nblast) (16, 58). The primary neurite tract was then manually removed from the neurons (by drawing an ROI), and the light-level tracing was compared to all the EM tracings from the AV1 tract by using the NBLAST algorithm (16, 58).

### Immunohistochemistry

Immunohistochemistry with antibodies was done similarly to (10), and the chemical labeling similarly to (6), with the exception of an overnight blocking step being used for antibody stainings. Primary antibodies used were: 1:20 mouse antinc82 (DSHB, University of Iowa, USA), 1:1600 chicken anti-GFP (ab13970, Abcam, Cambridge, UK), 1:200 rabbit anti-GABA (A2052, Sigma-Aldrich, MO, USA), and 1:400 mouse anti-ChAT (4B1, DSHB, University of Iowa, USA). Secondary antibodies were: Alexa-488 Goat anti-chicken, Alexa-568 Goat anti-Rabbit, Goat anti-mouse 633, all 1:800 (Life Technologies, Carlsbad, CA). For the chemical labeling, 1:1000 concentrations of SNAP-Surface 488 (NEB #S9124, New England Biolabs, Ipswich, MA), TMR Halo (G8252, Promega, Madison, WI) were used. Finally, brains were mounted on charged slides (Menzel-Glaeser, Braunschweig, Germany) using Vectashield (Vector Laboratories) as the mounting medium.

### Electrophysiology

*In vivo* patch-clamp recordings from the DA2 projection neurons were carried out as described in (17) using the R85E04 driver line and mCD8::GFP to label the neurons. Analysis of recordings used the open source gphys R (CRAN, http://www.r-project.org) package (see http://jefferis.github.io/gphys).

### *In vivo* calcium imaging

Functional imaging experiments on LHAV1a1 neurons were performed on flies containing two copies of UAS-GCaMP3 (at attP18 and attP40) driven by LH728 Split-GAL4 driver. GCaMP3 was used instead of newer versions of GCaMP for its higher baseline fluorescence which allowed easier identification of the neurons. Flies were placed into custom built holders, leaving the head and thorax exposed, under CO2 anaesthesia and secured in place with UV curable glue (Kemxert, KOA 300). Wax was used for securing the legs and the proboscis. A window was then cut into the head capsule with sharp forceps, and trachea and air sacks were removed in order to uncover the brain. Fly brains were bathed in external saline ([94]) adjusted to 275mM and 7.3 pH, and bubbled with 5% CO2. The saline had the following composition (Concentration, mM): NaCl 104.75; KCl 5; NaH2PO4 1; MgCl2.6H2O 1; CaCl2.2H2O 1; NaHCO3 26; TES 5; glucose 10; trehalose 10. The antennae were left under the holder so that they could be exposed to odour stimuli. A custom-built setup based on the Sutter (Novato, CA) Movable Objective Microscope with a Zeiss W Plan-Apochromat 20x/1.0 objective was used for the two photon imaging. A Coherent (Santa Clara, CA) Chameleon Vision Ti-Sapphire provided excitation, and image acquisition was controlled by ScanImage software (63). Image acquisition and odour delivery were triggered by a separate PC via Igor Pro software (Wavemetrics, Lake Oswego, OR) running Neuromatic. Images were captured at 8Hz at 265×255 pixel, and two photon excitation was provided at 900 nm. Odour stimulation was performed largely similarly to (24). Odour delivery started at 3000 ms after the beginning of a trial and lasted for 2000 ms. Image analysis was performed with custom scripts written in R employing the open source scanimage package (see https://github.com/jefferis/scanimage, 10.5281/zenodo.1401028). Data was both manually checked for motion artifacts, and excluded from the analysis if there were larger than 5% dF/F peaks during the baseline recording epoch, or if there were not larger than 5% dF/F responses to any of the tested odours during the stimulation epoch.

### Statistical analysis

All statistical analysis was performed in R (https://cran.r-project.org/). Shapiro-Wilk or Kolmogorov-Smirnov tests were used for assessing normality of distributions. Normally distributed data was then analysed by using One or Two-way ANOVAs, Welch’s one or two-sample t-tests, or Paired samples t-tests, whereas non-normally distributed data with Kruskal-Wallis rank sum tests, Wilcoxon rank sum tests, and Wilcoxon signed rank tests, where appropriate (see also Supplementary Table 1). For the egg-laying two-choice behavioural experiments (with R85E04 and LH728), power testing was done with effect size estimates based on data obtained on preliminary data on wild type and anosmic mutants (Ir8a1; Ir25a2; Orco1, Gr63a1) (values used for power test estimates were wild type mean PI=-.45, SD=.6; anosmic mean PI=0, SD=.6, which gives an effect size estimate of Cohen’s d=0.75), and based on this the required sample size for the experiments was 38.66, taking into account the Bonferroni corrected significance values. The power size estimations were done with the R package pwr (https://cran.r-project.org/web/packages/pwr).

The randomization tests for comparing DA2 downstream target morphology (see also ‘Clustering of neuron morphology’ below) to the number of DA2 inputs (Figure S2F) was conducted with custom R scripts. First, the coefficient of variation (CV) of DA2 inputs for each morphological cluster was calculated as

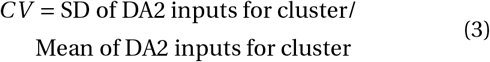

The DA2 input numbers were then randomly reassigned to clusters repeatedly (n=10.000), and new CVs were calculated for each iteration. Finally, the weighted mean (weighted by the proportion of neurons belonging to the cluster) of the original CVs were compared to the distribution of mean CV values obtained by the reshuffling. This was not done on ‘other’ neurons as only a few morphological clusters had multiple neurons in them, which makes it impossible to assess the variability of a cluster.

The randomization tests for aversion index (AI) values of completed LHNs were conducted with custom R scripts. First, the connectivity matrix of all excitatory uPNs (with known valence) was obtained via the R package catmaid (https://github.com/natverse/rcatmaid). In the cases of multiple sister PNs innervating the same glomerulus, the PNs were collapsed together into a PN type by taking the sum of their connections. All PN types were then assigned a valence (“aversive” or “not aversive”) based on earlier literature. After removing PNs of unknown valence, AI values for the LHNs were calculated as the ratio of the sum of connections from aversive PNs and the total sum of connections from excitatory uPNs. For the randomisation tests, DA2 PNs were excluded from the sample to avoid bias, and new (non-DA2) AI values were calculated. The valence labels were then randomly reshuffled (n=1000) and the AI values recalculated. The number of aversive and not aversive PN types was held constant, and the same as for the original data throughout. For each LHN, the observed distribution of random AI values was then compared to the original value, and the neuron was considered to significantly integrate aversive input if the observed AI value was higher than 95% of the values obtained by reshuffling.

Behavioural and imaging data throughout the paper are presented as notched box plots. The box represents the interquartile range of the sample (IQR, 25th - 75th percentiles) and is split by the median line. The whiskers extend to 1.5 × IQR beyond the box and the notches represent the 95% confidence interval for the sample median. The points mark individual sample points and asymmetrical notches indicate skewed distributions.

### Data availability

The FAFB reconstructed neurons will be shared with the Virtual Fly Brain project upon publication and will be available from (https://fafb.catmaid.virtualflybrain.org/). In addition, we are providing the skeletons as SWC files and a connectivity matrix as supplementary files.

