## Supplemental Materials for "Neural circuit basis of aversive odour processing in *Drosophila* from sensory input to descending output"

### Supplementary Information

#### Supplementary Table 1: Key resource information for figures

| Figure panel | Additional information |
|---|---|
| Figure 1A | A JFRC2 registered maximum intensity projection of a female brain. Raw image data from FlyLight database (http://flweb.janelia.org/cgi-bin/flew.cgi) (5). |
| Figure 1B | DA2 PNs are silenced by expressing tetanus toxin light-chain (TNT) (7) via R85E04. Groups are compared by a Kruskal-Wallis rank sum test followed by planned contrast tests of R85E04/TNT+ to each of the parental groups by Wilcoxon rank sum tests with Holm-Bonferroni corrections. MB247-GAL4/TNT+ was compared to 0 with a Wilcoxon signed rank test with continuity correction. |
| Figure 1C | Projections neurons traced from confocal image stacks. Raw data for DA2, DL4, V and DA4l PNs from Flycircuit (15, 16). DL5 PN originally traced for (10). |
| Figure 1G | Neurons are shown in the FAFB template brain. |
| Figure 3A | Mask generated from confocal images of DA2 PN axons in R85E04 registered onto the JFRC2 template brain. |
| Figure 3B | A maximum intensity projection of a R22D02 confocal stack registered onto the JFRC2 template brain. |
| Figure 3C | The PN mask together with R22D02 registered onto the JFRC2 template brain. A partial projection. |
| Figure 3E | Two split-GAL4 lines (LH728 and LH1983) are compared to the parental control Empty-Split GAL4 by two-sample t-tests with Holm-Bonferroni corrections for multiple comparisons. Gr66a-GAL4 shown as a positive aversive control. |
| Figure 3F-G | Maximum intensity projections of female brains registered onto the JFRC2013 template brain. |
| Figure 3H | Bridging registrations were used to transform the EM skeletons into the JFRC2013 reference brain by using nat.flybrains package for R. Visualised in Amira. |
| Figure 3K | LHAV1a1 LHNs are silenced by expressing tetanus toxin light-chain (TNT) (7) via LH728. Groups are compared by a Kruskal-Wallis rank sum test followed by planned contrast tests of LH728/TNT+ to each of the parental groups by Wilcoxon rank sum tests with Holm-Bonferroni corrections. |
| Figure 3L | LHAV1a1 LHNs are silenced by expressing tetanus toxin light-chain (TNT) (7) via LH1983. Groups are compared by Welch's Two Sample t-test with Holm-Bonferroni corrections. |
| Figure 3M | Groups are compared by a Wilcoxon rank sum test with continuity correction. |
| Figure 3N | GCaMP3.0 expressed via LH728. Area under curve compared by a paired samples t-test. |
| Figure 4B-C | Segmented maximum intensity projections registered onto the JFRC2013 reference brain. |
| Figure 4D | Bridging registrations were used to transform the EM skeleton into the JFRC2013 reference brain. Visualised in Amira. |
| Figure 4E | DNp42 neurons silenced by expressing tetanus toxin light-chain (TNT) (7) via SS57807. Planned contrasts done with Wilcoxon rank sum tests. |
| Figure 4F | Fly velocities measured in the FlyPEZ assay. Data was analysed by a two-way ANOVA to look for an interaction between the retinal (+/-) and genotype, followed by planned contrast tests with Wilcoxon rank sum tests. |

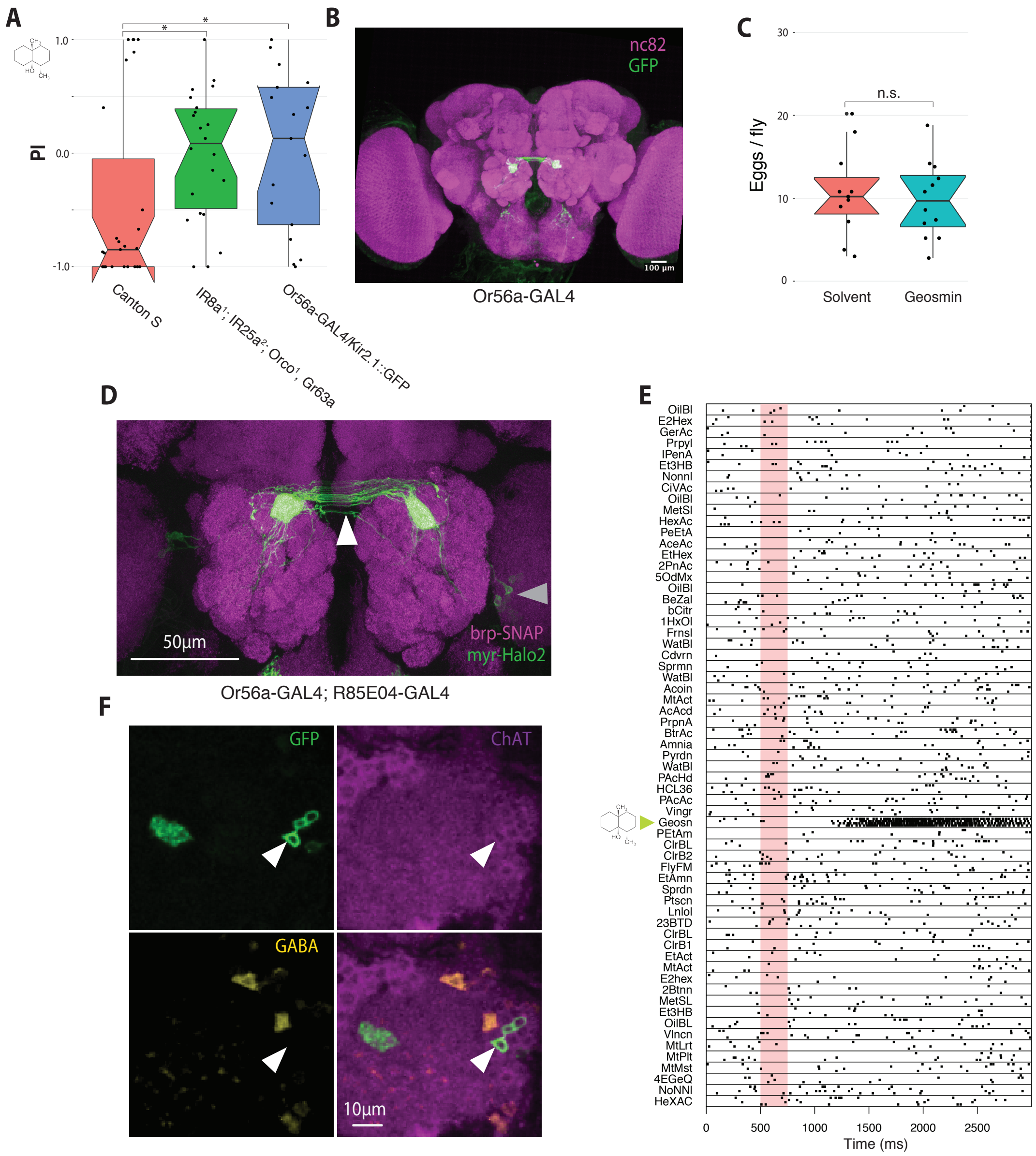


**Figure S1:** Wild-type flies avoid geosmin, which is sensed through Or56a ORNs that synapse onto the extremely narrowly tuned cholinergic DA2 PNs. (A) Egg-laying two-choice preference indices (PI) to geosmin for wild-type (Canton S) flies, anosmic mutants, and flies with silenced geosmin responsive sensory neurons (Or56a ORNs). The chemical structure for geosmin marks the stimulus side. Groups are compared by a Kruskal-Wallis rank sum test followed by planned comparisons of wild type group to the other groups by Wilcoxon rank sum tests with Holm-Bonferroni corrections. n=27 (left), n=22 (middle), and n=17 (right). (B) A maximum intensity projection of a Or56a-GAL4 confocal stack. Green channel marks the GFP, and magenta the nc82 neuropil stain. (C) Eggs laid per wild type fly under geosmin or solvent exposure (Welch's Two Sample t-test, n=12, n=11). (D) A high-resolution close-up of the antennal lobe expression pattern of Or56a-GAL4; R85E04. A maximum intensity projection of a female brain. Green channels marks the driver line expression pattern (Halo2 tag) and magenta the neuropil (SNAP tag). The arrowheads mark the midline crossing typical of ORN axons (white), and the PN cell bodies (grey). (E) Raster plot of *in vivo* patch clamp recordings from DA2 PNs (R85E04-GAL4) to a panel of odorants (17). Odour valve opening denoted by light red bar, timebase in ms. (F) Immunostainings against GFP, ChAT, and GABA in R85E04-GAL4, and the merge of the three. Single slices of confocal stacks showing the AL (magenta counterstain) and DA2 PN cell bodies (arrowheads). Significance values: * $p<0.05$ (G) Comparison of axo-axonic inputs from DA2 PNs to all PNs in the FAFB and Hemibrain EM volumes. PNs are coloured by valence.

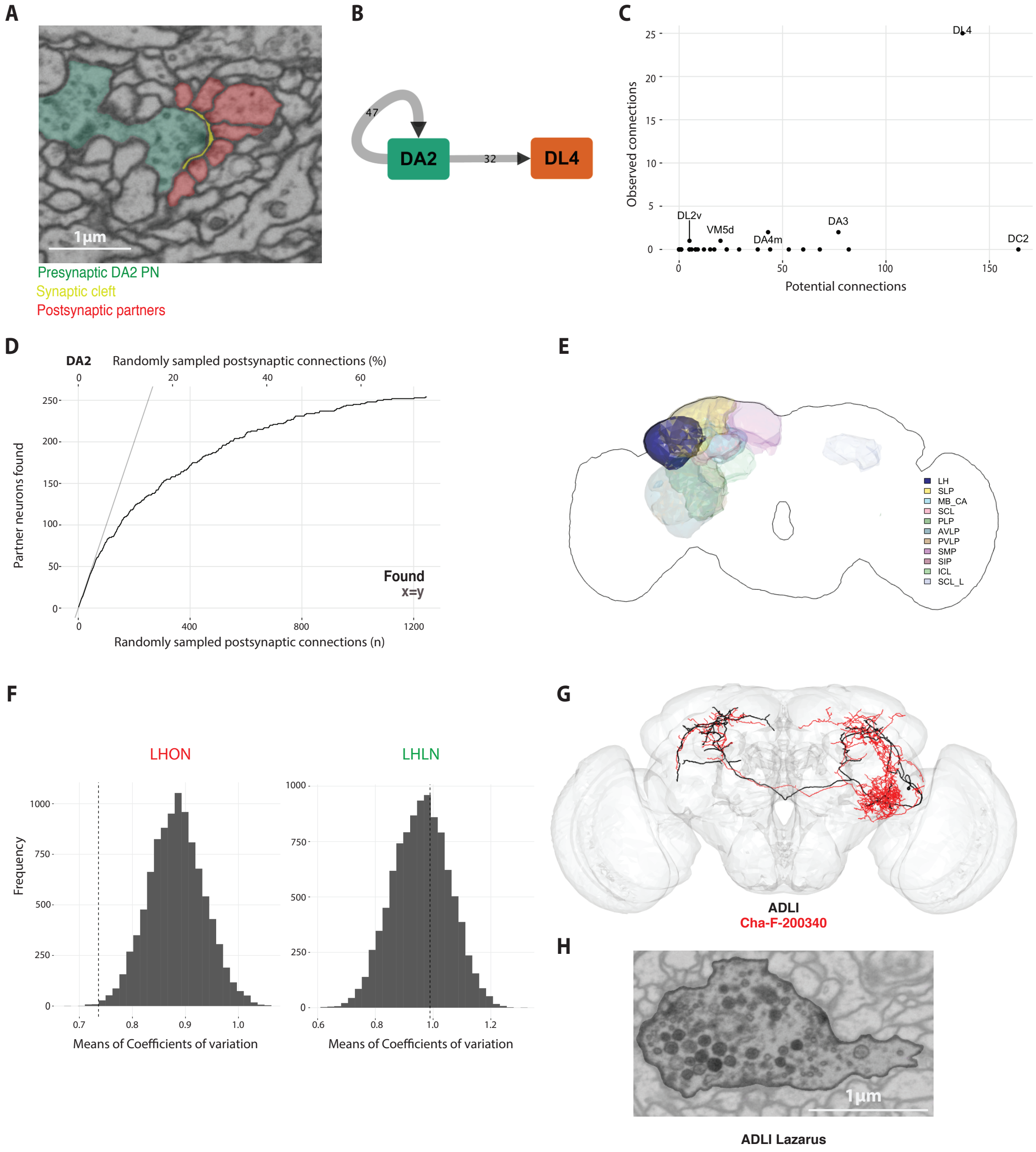


**Figure S2:** Supplementary information for DA2 downstream tracing. (A) An example of a synapse in the FAFB whole-brain EM volume. DA2 PN is highlighted in green, with the presynapse visible, synaptic cleft in yellow, and the profiles of 7 different postsynaptic neurons in red. (B) A connectivity diagram of axo-axonic PN synapses between DA2 PNs and the DL4 PN on the left hemisphere. (C) Observed versus estimated potential synapses from DA2 PNs to other nearby PNs in the LH. (D) The completion curve for the random sampling from the completed DA2 PN. Top x axis shows the % of postsynaptic connections sampled (of all identified synapses in the neuron), bottom x axis the number. Black line represents the number of identified postsynaptic partners (neuron fragments that could not be traced to a soma are excluded) as a function of the number of synaptic connections sampled. Grey line is x=y. (E) Neuropil regions targeted by LHONs downstream of DA2 PN. Colour intensity is based on the number of presynaptic connections in that neuropil. LH=lateral horn, SLP=superior lateral protocerebrum, AVLP=anterior ventrolateral protocerebrum, SCL= superior clamp, PLP=posteriorlateral protocerebrum, PVLP=posterior ventrolateral protocerebrum, SMP=superior medial protocerebrum, SIP=superior intermediate protocerebrum, MB_CA=mushroom body calyx, ICL=inferior clamp. (F) Observed (dotted line) versus randomised cluster variation (histogram) in DA2 input for LHONs and LHLNs. Coefficients of variation of DA2 input (cluster SD/cluster mean) were calculated for each anatomical cluster, and the mean of these was taken to represent the within-cluster variation in DA2 input. DA2 inputs were then randomly reassigned to clusters repeatedly (n=10.000), and the resultant distribution of means of coefficient of variation compared to the original value. For LHONs 99.8% of the reshuffled iterations had a higher mean coefficient of variation than the observed original value. For LHLNs, the same was true only for 38%. (G) EM traced ADLI neuron (black) overlaid with its Flycircuit (15, 16) match Cha-F-200340 (red) in the FCWB reference brain. (H) An example EM image showing dense core vesicles in ADLI.

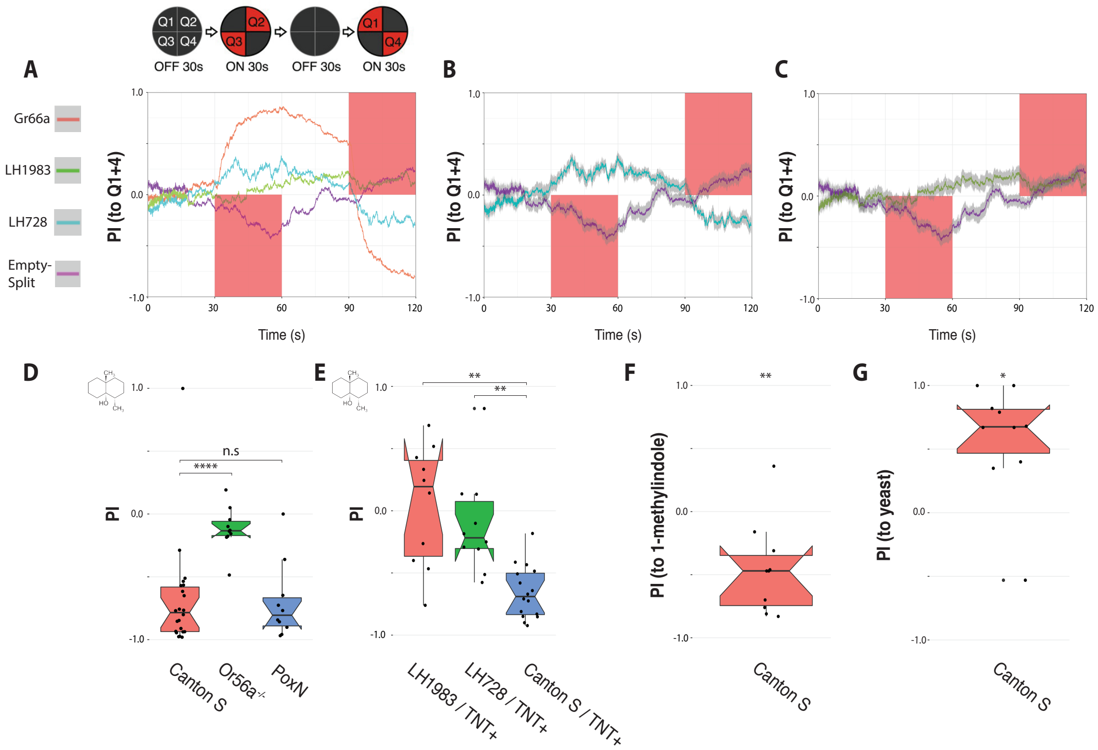


**Figure S3:** Supplementary information for behavioral experiments with LHAV1a1 neurons. (A) The mean PI (to Q1+Q4) as a function of time for LH728, LH1983, Empty-Split GAL4 (genetic control), and Gr66a-GAL4 (a positive aversive control) in the optogenetic behavioural assay. The red squares mark the stimulation epochs. n=16 for all genotypes. (B) The mean PI and SEM of LH728 plotted against the control. (C) The mean PI and SEM of LH1983 plotted against the control. (D) Egg-laying two-choice PI to geosmin for wild-type, Or56a, and poxn mutants in the split-plate assay. Groups are compared by a Kruskal-Wallis rank sum test, followed by planned comparisons with Wilcoxon rank sum test with Holm-Bonferroni corrections. n=22 (left), n=10 (middle), n=10 (right). (E) Egg-laying two-choice PI to geosmin while silencing LHAV1a1 neurons in the split-plate assay. Groups are compared with a one-way ANOVA, followed by planned comparisons of the L1983/TNT+ and L728/TNT+ groups to the parental control group by t-tests with Holm-Bonferroni corrections. n=10 (left), n=10 (middle), n=16 (right). (F) Egg-laying two-choice PI to 1-methylindole for wild-type flies (n=10) in the split-plate assay. The PIs were compared to 0 by using a One-sample t-test. (G) Egg-laying two-choice PI to yeast odor for wild-type flies (n=10). The PIs were compared to 0 by using a One-sample t-test. Significance values: * $p<0.05$ ** $p<0.01$ *** $p<0.001$ **** $p<0.0001$

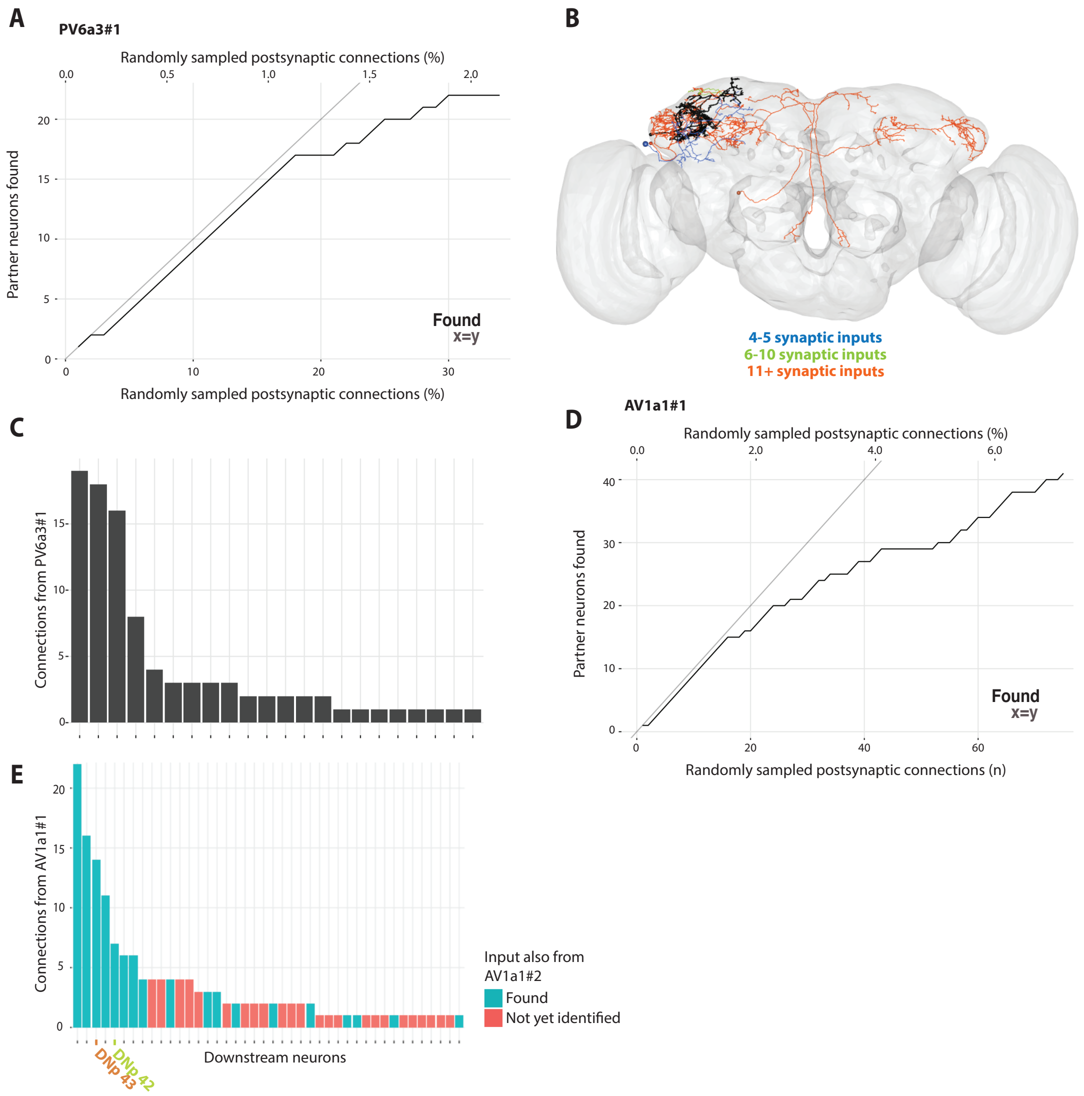


**Figure S4:** Supplementary information for downstream tracing from PV6a3#1 and LHAV1a1 neurons. (A) The completion curve for the random sampling from PV6a3#1. Top x axis shows the % of presynaptic connections sampled (of all identified synapses in the neuron), bottom x axis the number. Black line represents the number of identified postsynaptic partners (neuron fragments without a soma are excluded) as a function of the number of synaptic connections sampled, and grey line is x=y. (B) Identified downstream targets of PV6a3#1 with 4 or more synaptic inputs. Neurons are colour-coded based on connection strength. (C) The number of synaptic inputs from PV6a3#1 to each identified downstream target. (D) The completion curve for the random sampling from the LHAV1a1#1. Top x axis shows the % of synapses sampled (of all identified synapses in the neuron), bottom x axis the number. Black line represents the number of identified postsynaptic partners (neuron fragments without a soma are excluded) as a function of the number of synaptic connections sampled, and grey line is x=y. (E) The number of synaptic inputs from LHAV1a1#1 to each downstream target. The DNs are highlighted. The neurons that also receive input from LHAV1a1#2 are marked in cyan.

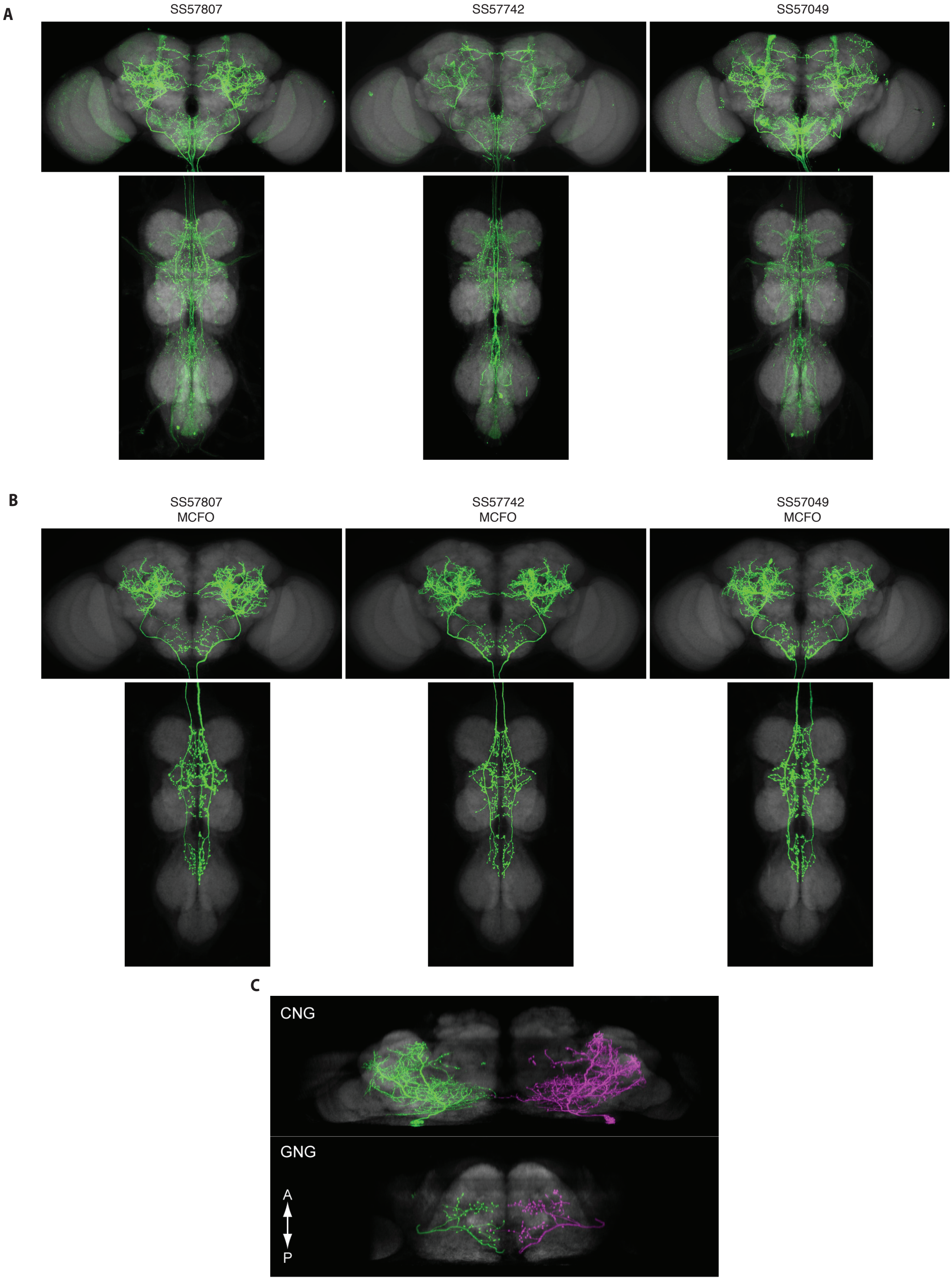


**Figure S5:** Confocal images for DNp42 driver lines. (A) Maximum intensity projections of confocal stacks of Split-GAL4 driver lines labeling DNp42. Green channel marks the GFP, and grey the nc82 neuropil stain. (B) Maximum intensity projections of confocal stacks of MCFOs of Split-GAL4 driver lines labeling DNp42. Green channel marks the GFP, and grey the nc82 neuropil stain. (C) Dorsal (top) and ventral (bottom) views of maximum intensity projections of MCFO labeled DNp42 neurons.

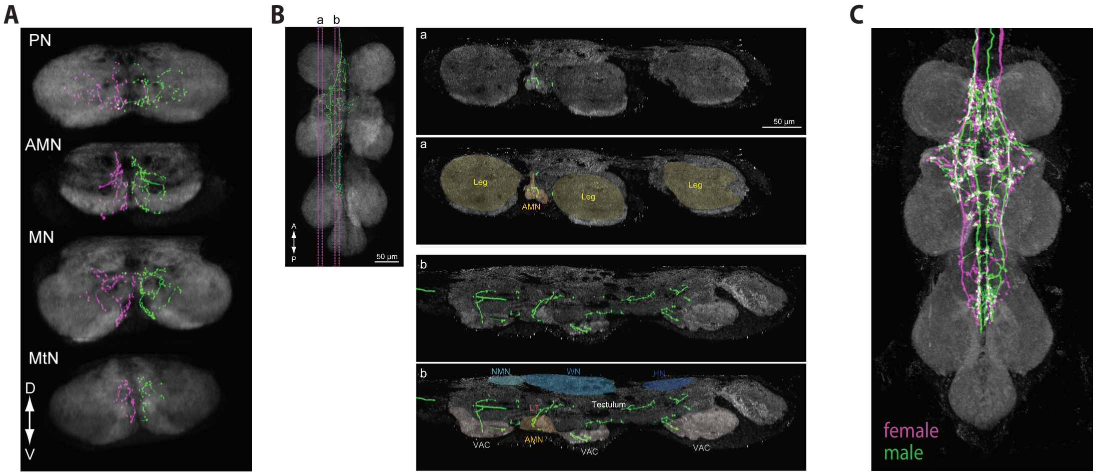


**Figure S6:** DNp42 projections in the VNC. (A) Transverse sections of registered MCFO images of DNp42 neurons in the VNC at different planes. (B) Sagittal sections of the registered MCFO images of DNp42 neurons in the VNC at two different planes (a and b). Left panel shows the location of planes. The top panels for a and b show the expression pattern, whereas the bottom panels show the expression together with labeled neuropil areas. (C) A comparison of female and male DNp42 neurons in the VNC. ProNm=prothoracic neuromere, AMNn=accessory mesothoracic neuropil, MesoNm=mesothoracic neuromere, MetaNm=metathoracic neuromere, Leg=leg neuropil, NTct=neck tectulum, WTct=wing tectulum, HTct=haltere neuropil, VAC=ventral association center, LTct=lower tectulum.
